# Unearthing the Boreal Soil Resistome Associated with Permafrost Thaw

**DOI:** 10.1101/2020.10.19.346478

**Authors:** Tracie J. Haan, Devin M. Drown

## Abstract

Understanding the distribution and mobility of antibiotic resistance genes (ARGs) in soil bacteria from diverse ecological niches is critical in assessing their impacts on the global spread of antibiotic resistance. In permafrost associated soils, climate and human driven forces augment near-surface thaw altering the overlying active layer. Physiochemical changes shift bacterial community composition and metabolic functioning, however, it is unknown if permafrost thaw will affect ARGs comprising the boreal soil resistome. To assess how thaw shifts the resistome, we performed susceptibility testing and whole genome sequencing on soil isolates from a disturbance-induced thaw gradient in Interior Alaska. We found resistance was widespread in the Alaskan isolates, with 87% of the 90 isolates resistant to at least one of the five antibiotics. We also observed positive trends in both the proportion of resistant isolates and the abundance of ARGs with permafrost thaw. However, the number of ARGs per genome and types of genes present were shown to cluster more strongly by bacterial taxa rather than thaw emphasizing the evolutionary origins of resistance and the role vertical gene transfer has in shaping the predominantly chromosomally encoded ARGs. The observed higher proportion of plasmid-borne and distinct ARGs in our isolates compared to RefSoil+ suggests local conditions affect the composition of the resistome along with selection for ARG mobility. Overall taxonomy and geography shape the resistome, suggesting that as microbial communities shift in response to permafrost thaw so will the ARGs in the boreal active layer.

**IMPORTANCE:** As antibiotic resistance continues to emerge and rapidly spread in clinical settings, it is imperative to generate studies that build insight into the ecology of environmental resistance genes that pose a threat to human health. This study provides insight into the occurrence of diverse ARGs found in Alaskan soil bacteria which is suggestive of the potential to compromise health. The observed differences in ARG abundance with increasing permafrost thaw suggest the role of soil disturbance in driving the distribution of resistant determinants and the predominant taxa that shape the resistome. Moreover, the high-quality whole genome assemblies generated in this study are an extensive resource for microbial researchers interested in permafrost thaw and will provide a steppingstone for future research into ARG mobility and transmission risks.

## INTRODUCTION

The rapid evolution and spread of antibiotic resistance is one of the greatest challenges faced in public health today. Antibiotic resistance impedes the successful treatment of bacterial infections by reducing antibiotic efficacy, increasing disease burden, mortality rates, hospitalization time and cost (O’Neill 2016). In 2017 alone, the United States was estimated to have 2,868,700 antibiotic-resistant infections resulting in 35,900 deaths (Center for Disease Control and Prevention 2019). On an evolutionary time scale, the extensive prevalence of resistant phenotypes in human pathogens is a recent event driven by the large-scale production and widespread use of antibiotics in clinical, agricultural, and veterinary settings (Ghosh et al. 2007, He et al. 2020). Even when antibiotic stewardship is instilled (i.e. antibiotic use is confined to essential needs) antibiotics, or pollutants such as heavy metals that co-select for resistance (Berg et al. 2005), are dispersed within microbial habitats thereby generating selective pressures that may increase the abundance of resistant strains and their associated antibiotic resistance genes (ARGs). It was originally thought genetic variability driving resistance was primarily caused by mutational modification to antibiotic targets, and thus would remain clonal (Djordjevic et al. 2013). However, it is now evident that mutational driven resistance is a weaker force compared to ARGs acquired via horizontal gene transfer (Aminov 2009). In pathogens, resistance genes can be acquired from diverse microbial habitats and taxa (Djordjevic et al. 2013), including bacteria from pristine environments free of antibiotics introduced via human activities (D’Cost et al. 2011). It is therefore important to assess which bacterial taxa in distinct microbial biospheres are the predominant contributors to the evolution of resistance in pathogens (Forsberg et al., 2012).

Soils, one of the most diverse microbial habitats on earth, are a vast repository of both antibiotic producing and coevolved resistant microbial taxa. Antibiotic production is thought to have originated in soils from 2 Gyr to 40 Myr ago suggesting resistance had a concomitant evolution over a similar timeframe (Baltz 2008; Hall et al. 2004). The evolutionary origins of resistance is also supported by studies that have unveiled soil environments unpolluted by human activity, such as 30,000-year-old Beringian permafrost sediments (D’Costa et al. 2011), that harbor diverse resistance mechanisms to modern antibiotics. Moreover, bacteria such as *Streptomyces*, an Actinomycete genus that produces around two-thirds of clinically used antibiotics, are abundant in soils (de Lima Procópio et al. 2012, Watve et al. 2001). The presence of these antibiotic-producing genera thus promotes the evolution, and potential dissemination via horizontal gene transfer, of clinically relevant resistance genes. Recent studies have reported ARGs in soil-borne bacteria identical to those circulating in pathogenic isolates which is suggestive of horizontal gene transfer events having occurred (Finley et al. 2013, Forsberg et al. 2014). These shared ARGs between soil-borne and pathogenic bacteria emphasize the potential role soils have in the evolution and dissemination of resistance globally. In order to better assess risks posed by soil-borne ARGs, more attention should be paid to environmental factors that favor growth of resistant bacteria, as well as conditions that promote ARG selection and mobility such as plasmid carriage (Aminov and Mackie, 2007). Revealing the ARG distribution and conditions that affect antibiotic resistance in distinct ecosystems, especially those affected by climactic or human driven changes, can help us understand the processes that lead to the emergence of antibiotic - resistant pathogens.

Alaskan soils are one environment undergoing unprecedented change as warming within the arctic occurs 2.5 times faster than the rest of the globe triggering a rise in the frequency of soil disturbance events, such as wildfires and thermokarst formation (Schuur et al. 2018). These disturbances are of particular concern in Alaska because approximately 85% of land is underlain by discontinuous permafrost (McGuire et al. 2018). With both climatic and human-driven soil disturbances permafrost, the frozen amalgam of rock, soil, and ice that remains below freezing for at least two years, experiences augmented thaw. Permafrost thaw subsequently affects the active layer soils that overlay permafrost by shifting the physical and chemical properties (Douglas et al. 2008). Since active layer soils defines nutrient availability, rooting space for plants, and microbial respiration, thaw driven changes to it affect arctic ecosystems at large. On a smaller scale, alterations to pH, moisture, and nutrients in the active layer allow microbial communities to gain access to large pools of previously frozen stores of carbon, which not only affects ecosystem function, but shifts microbial community composition (Schurr et al. 2007). These changes reveal new niches allowing bacterial variants that are better suited for colonization of alternative niches to thrive. Thus the shift in community composition is likely driven by selective forces that favor microbial taxa capable of gaining access to newly revealed soil resources or that carry defense mechanisms allowing bacterial cells to cope with environmental stressors induced by thaw, such as the as release of biogenic volatile organic compounds (BVOC) (Kramshøj et al. 2018).

Permafrost thaw may directly impact the conditions that favor the selection of antibiotic resistant bacterial taxa. For example, shifting active layer conditions may favor bacterial species that are capable of competitive inhibition via antibiotic production therefore directly selecting for bacteria resistant to antibiotics generated by these producers. Shifting conditions could also select indirectly for mechanisms that allow bacteria to concurrently cope with environmental stressors and antibiotics, such as efflux pumps (Hibbing et al. 2010). The prevalence of antibiotic producing bacterial taxa may also have additional ecological impacts, such as impairing ecosystem function and service as a result of community changes driven by antibiotic production (Ashbolt et al. 2013). Thus, a shift in the abundance of antibiotic producers in a community may be a concern for reasons other than the obvious role in promoting selection for resistance. Moreover, since plasmid-mediated genetic variation allows bacterial populations to respond rapidly to environmental challenges, permafrost thaw may therefore increase the abundance of mobile genetic elements, such as plasmids (Djordjevic et al. 2013) and integrons (Stalder et al. 2012), and thus the abundance of ARGs housed on plasmids that pose more risk in terms of dissemination to pathogenic bacteria (Aminov et al. 2009).

The goal of this study is to assess how permafrost driven shifts will affect the abundance and mobility of ARGs that compose the Alaskan active layer resistome. To address this goal we identified ARGs in the whole genome sequences of bacteria cultured from active layer soils from across a permafrost thaw gradient in Fairbanks, Alaska. We compared ARGs across this thaw gradient to those identified in bacteria from database of global soil bacteria, RefSoil+ (Dunivin et al. 2019). This larger database of soil bacteria containing both the whole genomes and plasmids sequences from cultured soil bacteria of global soil habitats makes it possible to investigate distinguishing features of ARG ecology in Alaskan isolates, such as the ARG plasmid carriage. Overall these methods will allow us to gain insights into both the role permafrost thaw has in shaping the active layer soil resistome along with shedding light on the role of horizontal gene transfer has in the soil communities evolution of antibiotic resistance that may influence the resistance in pathogenic bacteria.

## METHODS

### Permafrost Thaw Gradient

The Fairbanks Permafrost Experiment Station (FPES) is an ice-rich permafrost site in Interior Alaska (64.875646°N, 147.668981°W) established by the Army Corps of Engineers as part of the Cold Regions Research and Engineering Lab. The site consists of three 3,721 m^2^ Linell plots (Linell 1975) with increasing levels of disturbance now known as the Un-Disturbed (UD), Semi-Disturbed (SD), and Most-Disturbed (MD) sites (Figure 1). In 1946 the three sites were established to simulate soil disturbance events, such as wildfire or anthropogenic disturbance, on permafrost degradation by clearing of vegetation. The UD site was left undisturbed to preserve the subarctic taiga forest, whereas the SD site had the surface vegetation cleared while roots and soil organic matter were left intact and the MD site had both surface vegetation and organic matter removed. In 2007 the UD site, which is now monitored as part of the Circumpolar Active Layer Monitoring Network (CALM), was found to have little to no thaw since 1946 whereas the SD and MD had up to 4.7m and 9.8m respectively (Douglas et al. 2008). These results suggested that permafrost degradation is dependent on both time and surface vegetation which has major implications when it comes to increasing frequency of disturbance events, like wildfires and thermokarst formation, influenced by climate change.

**Figure 1.**
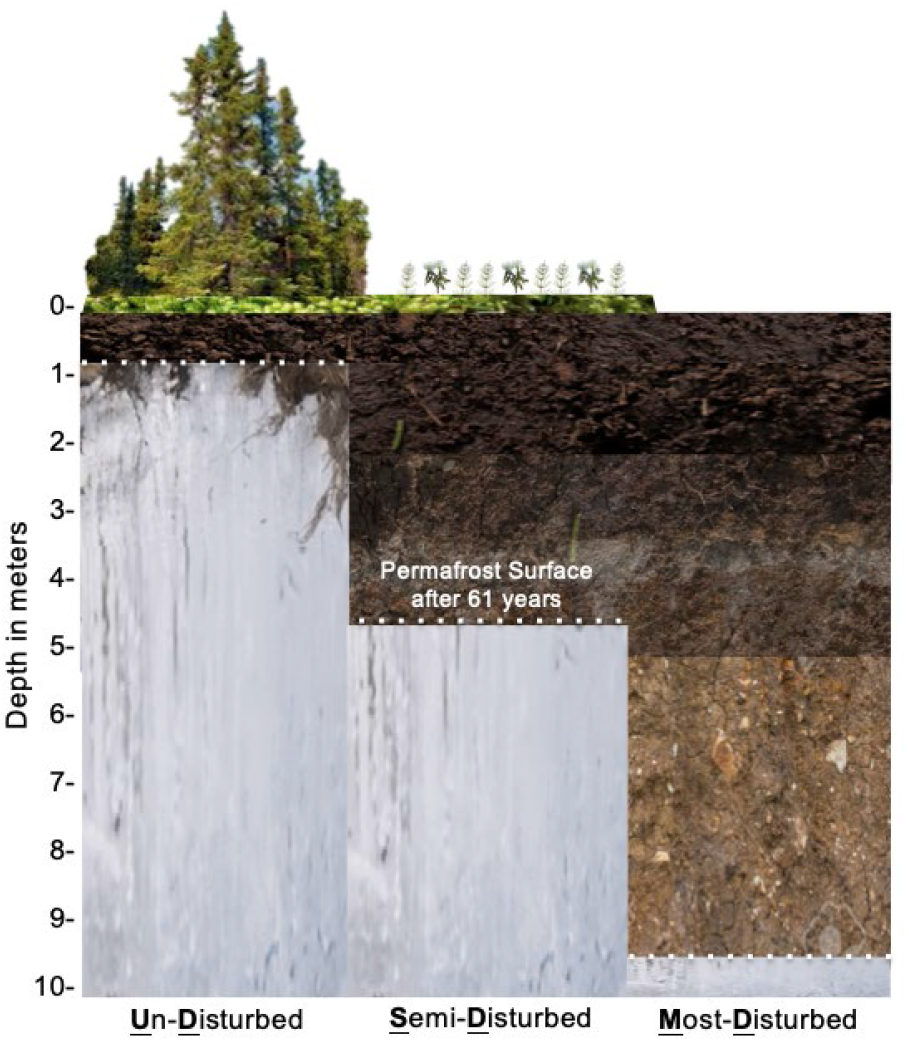
Diagram of the disturbance level sites (Undisturbed=UD, Semi-Disturbed=SD, Most-Disturbed=MD) at the Fairbanks Permafrost Experiment Station (FPES) and the subsequent depth of permafrost thaw after 61years.

Vegetation at the FPES is typical of the Alaskan Interior-subarctic taiga forest. The undisturbed site is a relatively open black spruce stand (*Picea mariana*) with an understory of continuous thick moss layer interspersed with low-bush cranberry (*Vaccinium vitis-idaea*) and Labrador tea (*Rhododendron groenlandicum*). The UD site can be classified as mesic (Johnstone et al. 2008) with a soil organic layer thickness ranging 2 to 35 cm thick, with little to no thaw during maximal permafrost thaw. The semi-disturbed site is now a mix stand dominated by black spruce, Alaskan paper birch (*Betula neoalaskana*), and willow (*Salix alaxensis*). The understory contains a mixture of Labrador tea (*Rhododendron groenlandicum*), Peltigera lichen, roses, horsetail, cloudberry, and small amounts of grass with little litter cover. The MD site is an open shrub land dominated by willows (*Salix alaxensis*) and a developing over story up to 5 m tall of Alaskan birch and black spruce (*Picea mariana*). The understory contains many grasses, clovers, horse-tail (*Equisetum*), and some bare ground. There is no permafrost within the top 4.7 m in either disturbed sites (Douglas et al. 2008).

### Bacterial Culturing

In the September of 2017 we collected two 10 cm wide by 20 cm deep soil cores from each FPES site with a sterilized corer. Prior to coring the top layer of moss and vegetation were removed. Soil cores were then extracted and immediately stored in a cooler throughout sample collection. Soil cores were moved to a +4°C fridge until processing 24 h later. In order to process the cores in a way that prevented contamination from any exogenous cells on the exterior of the soil core, outer portion of each core was removed using a sterile scalpel. The interiors of each core were then sub-sample using sterile forceps along a depth gradient at intervals of about 2.5 cm for 20 cm to generate a total of 1 g of soil. The 1 g of soil was used to inoculate 100ml tryptic soy broth (TSB) to produce an enrichment culture. After 48 h at 22°C, we plated serial dilutions (1:10, 1:100, 1:1000) of the enrichment culture three times for each sample and incubated the plates at three temperatures (+4°C, +12°C, and +20 °C), for a total of twelve plates per sample, until distinct colony formation was observed. Ten discrete colonies were chosen at random from each temperature and FPES site by using a random number generator to count colonies across transects moving across the plate horizontally from the top of the plate to the bottom. Only discrete colonies (i.e. colonies without overlapping colony growth) were counted along the transect lines. This process yielded 90 total colonies, 30 per FPES site. Each colony was isolated and purified using three rounds of streak plate method.

### Antibiotic Susceptibility Testing

We screened each isolate for antibiotic resistance using the Kirby-Bauer disk diffusion method (Hudzicki 2009) against five antibiotics (tetracycline, erythromycin, kanamycin, chloramphenicol, and ampicillin) each representing a distinct antibiotic class. Breakpoints from US Clinical and Laboratory Standards Institute M100, 30th ed. (https://clsi.org/) established for *Enterobacterales* were used for *Serratia, Pantoea*, and *Erwinia* isolates, *Pseudomonas* spp. for *Pseudomonas* isolates, and *Enterococcus* spp. for *Bacillus* and *Exinguobacterium* to determine if an isolate was susceptible, intermediate, or resistant to each antibiotic tested. This means that isolates without breakpoints for one of the tested antibiotics, due to physiological characteristics that render that genus resistant such as in the case of *Pseudomonas* and Ampicillin, were removed from subsequent analysis.

### Whole Genome Sequencing, Assembly, and Taxonomic Classification

From each purified isolate, we inoculated a liquid culture of TSB, incubated it at 22°C overnight, and used 1.8 ml of this liquid culture to extract DNA using the DNeasy UltraClean microbial kit (Qiagen, Venlo, Netherlands) following manufacture protocols. We used a Nextera XT library (Illumina, San Diego, CA, USA) prepared by the Genomics Core Lab at the University of Alaska Fairbanks to sequence on an Illumina MiSeq platform using version 3 reagents. We trimmed adapters with TrimGalore version 0.5.0 (https://github.com/FelixKrueger/TrimGalore). For long reads, a combination of SQK-RBK004 and VSK-VSK002 library preparation (Oxford Nanopore Technology [ONT]) (Table S2) were used followed by sequencing on a MinION device (ONT) and an r9.4.1 flow cell (FLO-MIN106) for 48-72 h. We base called the raw data using Guppy v3.4.5 (ONT) specifying the high-accuracy model (-c dna_r9.4.1_450bps_hac.cfg) and default parameters. We de-multiplexed isolate samples using the guppy_barcoder function of Guppy with parameters to discard sequences with middle adapters (− detect_mid_strand_barcodes) and trim barcodes (−trim_barcodes). We used Filtlong v0.2.0 (https://github.com/rrwick/filtlong) to filter by length (≥50 bp; −min_length 50) and quality (Q) score (≥10; −min_mean_q 90). The unicycler_polish tool of Unicycler v0.4.8 (Wick et al. 2017) was used for genome polishing with the Illumina reads and the Flye assembly as inputs. For isolates TH26 and TH81, ONT long-read assemblies previously published in Haan et al. 2019 and Humphrey et al. 2019 respectively were used as inputs for unicycler_polish. These assemblies were annotated with RAST tool kit (RASTtk) in PATRIC v3.6.3 using the Genome Annotation Service (Brettin et al. 2015). 16S rRNA gene copies for each assembly were aligned using MAFFT v7.450 and consensus sequences were run through BLASTn version 2.10.0 against the NCBI 16S rRNA database. The top five hits (ranked by bit score) were then used to assign taxonomy at a genus level (Table 1).

**Table 1.** Number of isolates by genus and FPES site.

| Bacterial Taxa |  | FPES Site |  |  | Total |
| --- | --- | --- | --- | --- | --- |
| Phyla | Genera | UD | SD | MD |  |
| *Firmicute* | *Bacillus* | 14 | 11 | 5 | 30 |
| *Firmicute* | *Exiguobacterium* | 0 | 0 | 1 | 1 |
| *Proteobacteria* | *Erwinia* | 1 | 1 | 3 | 5 |
| *Proteobacteria* | *Pantoea* | 0 | 0 | 2 | 2 |
| *Proteobacteria* | *Pseudomonas* | 14 | 18 | 19 | 51 |
| *Proteobacteria* | *Serratia* | 1 | 0 | 0 | 1 |
| **Total** |  | 30 | 30 | 30 |  |

### Antibiotic Resistance Gene Identification

There are a variety of published antibiotic resistance gene databases used for annotation of resistance genes, one of the most widely used being the Comprehensive Antibiotic Resistance Database (CARD) (McArthur et al. 2013). The CARD database provides well-developed and extensive antibiotic resistance ontology (ARO) and monthly curation updates to include the most up to date ARG reference data which is exclusively derived from peer-reviewed publications validated by clinical or experimental data (Alcock et al. 2020). A 2016 study found that CARD was able to outperform other popular AR databases including ARDB, ResFinder, and CBMAR by correctly identifying down to a variant level for the all variants of the two genes tested, bla_VIM_ and bla_NDM_, and unlike any of the other databases was able to accurately identify the maximum number of resistance genes from the whole genome sequences of 3 strains of methicillin resistant *Staphylococcus aureus* (Xavier et al. 2006). In this study we annotated each whole genome assembly with CARD version 3.0.9 using command line tool Resistance Gene Identifier (RGI) version 5.1.0 Main specifying input type contig (-t contig) with default parameters for BLAST alignment (-a BLAST) and strict and perfect hits only by not choosing the include loose option. We chose to use the strict algorithm rather than the perfect or loose because it can detect previously unknown homologs using detection models with curated similarity cut-offs while still ensuring the detected variant is a functional resistance gene rather than spurious partial hits (Jia et al. 2006). Results from RGI were further quality controlled by removing any ARO hits defined as mutations or ARO hits with less than 50% coverage of the reference sequence unless cutoff on the edge of a contig in order to ensure hits were ARG homologs rather than spurious hits. To determine if a hit was located on a chromosome or plasmid, contigs containing hits were run through BLASTn (Zhang et al. 2000) in Geneious Prime version 2019.2.1 against bacteria (taxid=2) database. The top hit ranked by bit score was then used to determine if the contig was most similar to a known plasmid or chromosome sequence.

### RefSoil+ Comparison

RefSoil+ (Dunivin et al. 2019) genomes and plasmids were downloaded from NCBI using accession numbers available on the RefSoil+ github page (https://github.com/ShadeLab/RefSoil_plasmids). Each RefSoil+ sequence was then run through RGI following the same protocol as the FPES analysis of ARGs. Genomes belonging to the matching genera (*Bacillus*, *Erwinia*, *Exinguobacterium*, *Pantoea*, *Pseudomonas*, and *Serratia*) as FPES isolates were used for comparison of the RefSoil+ and FPES resistance genes. The genus *Pseudomonas* contains the largest and most diverse species with eight distinct phylogenomic groups, because the species *Pseudomonas aeruginosa* is distinctive from other phylogenomic groups within our samples *aeruginosa* genomes were removed from FPES vs RefSoil+ analysis.

### Data Analyses and Statistics

Tabular outputs from RGI were used to conduct statistical analyses in R Studio version 3.5.7. (R Development Core Team, 2020) and visualizations were generated in the R package ggplot2 version 3.2.1 (Wickham 2016). In order to determine if ARGs were influenced more strongly by FPES site or genus, we examined statistical differences in the mean number of ARGs per genome between genera and across thaw sites in FPES isolates and between correspondent genera from RefSoil+. We used non-parametric Wilcoxon signed rank test to compare means between groups to see if specific sites or taxa had a stronger influence on the number of resistance genes within a group; mean ARG abundance across all groups was compared using Kruskal-Wallis one-way analysis of variance. Dendrograms of Euclidean distance were created along with a heat map that highlighted core ARGs versus accessory (i.e. acquired) across groups. The heat map with dendrograms was created by normalizing each group (e.g. Site and Genus-Bacillus from UD) by number of isolates within that group and then scaling the normalized counts to produce a heatmap of z scores for each ARG. The heat map visualization was generated in R using the package pheatmap (version 1.0.12; Kolde, 2019).

## RESULTS

### Antibiotic Susceptibility

Widespread resistance was observed in the isolates screened with 87% of the 90 total isolates exhibiting resistance to at least one of the five antibiotics tested. Ampicillin had the highest proportion of resistant isolates across thaw sites (UD=76.4%; SD=83.3%; MD=81.8%) whereas Tetracycline had the lowest (UD=3.3%; SD=3.3%; MD=0.0%). We observed a positive trend in the number of resistant isolates with thaw for Ampicillin, Chloramphenicol, and Erythromycin that was also observed for intermediate resistance to Kanamycin and Tetracycline (Figure 2).

**Figure 2.**
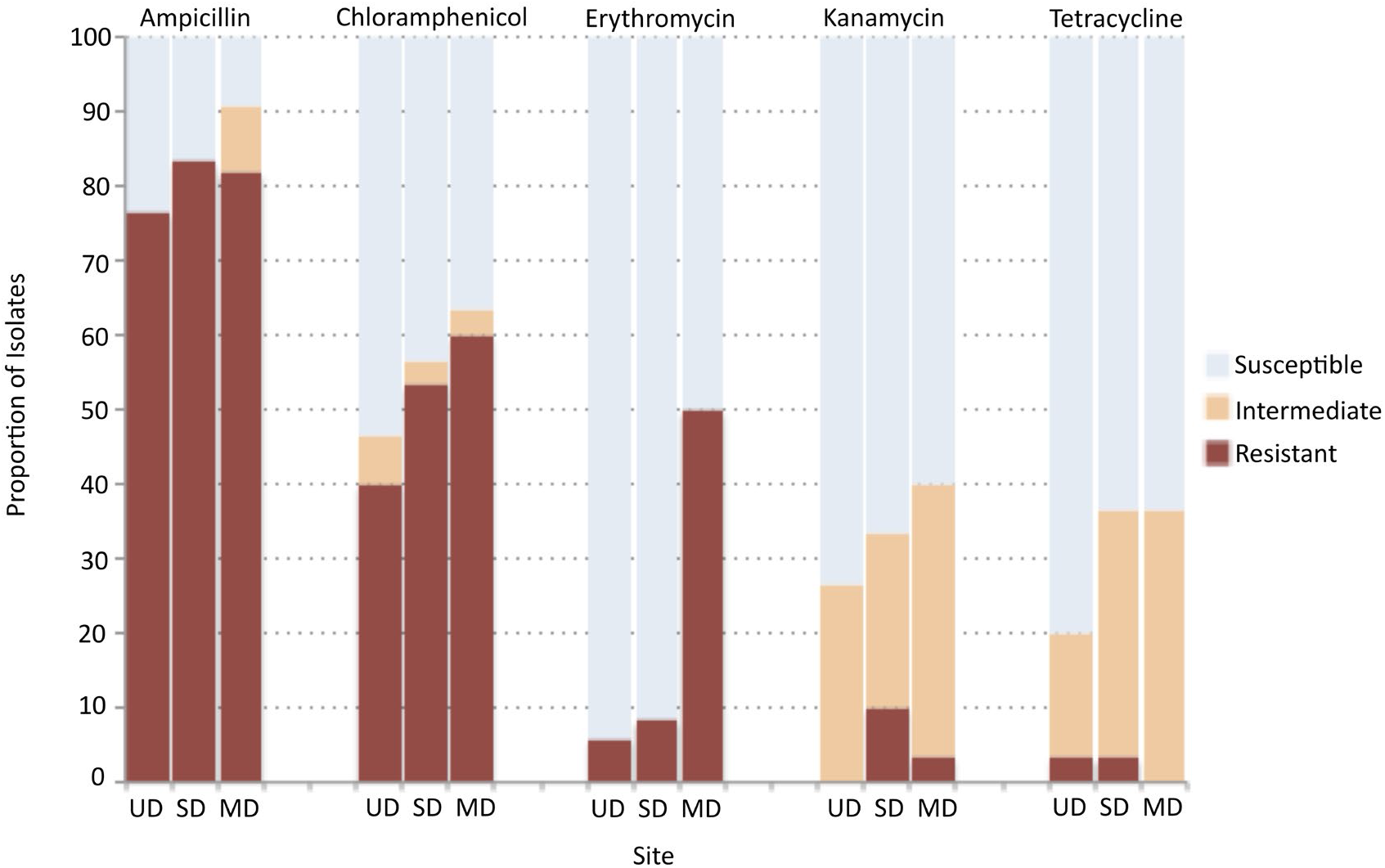
Proportion of isolates at each FPES thaw site and associated level of susceptibility to each antibiotic tested based on CLSI breakpoints.

### Genome Assemblies

The assembled genomes had a high mean percent completeness of 98.91 ± 0.50 and low percent contamination of 1.16 ± 0.26 suggesting high quality assemblies were produced (Table S3). The assemblies were also highly contiguous (N50 = 3,639,582 ± 279,870 bp, N50n = 13 ± 5 contigs) with mean total length of 6,096,842 ± 946,004 bp and mean GC content of 52 % ± 1 (Table S3).

### Antibiotic Resistance Genes

Across all FPES genomes RGI identified 379 significant hits comprising 27 CARD-based AROs (Table S1). Of these hits 30 had 100% sequence identity to CARD AROs and another 32 were highly similar (sequence identity >90%). Four genes hits, two encoding aminoglycoside inactivating enzymes AAC(6’)-32 and AAC(6’)-Ir and two encoding bcrC, an undecaprenyl pyrophosphate related protein, had full length coverage of the reference sequence along with 100% sequence identity. These findings show that some ARGs present in these soil isolates are highly homologous to those present in the CARD database rather than just ancient divergent homologs. However, overall mean percent identity across all hits identified was 68.4 % ± 19 suggesting that although some hits were highly similar to known resistance genes, there were also many hits identified that diverge from ARGs present in the CARD database suggesting novel homologs of known resistance genes are present in these isolates.

Genes from mechanisms of antibiotic resistance including antibiotic efflux, antibiotic inactivation, antibiotic target modification, and antibiotic target protection were observed. The top two most abundant resistance genes identified across isolates were genes encoding antibiotic efflux. The most abundant efflux pump was *adeF*, a gene encoding resistance-nodulation-cell division efflux pump that have been shown to confer multi-drug resistance when overexpressed in clinical *Acinetobacter baumannii* and *Pseudomonas aeruginosa* strains (Mobasseri et al. 2018). There were 176 chromosomally encoded gene copies distributed across the 100% of *Proteobacteria* isolates with a high coverage of the gene across assemblies (% length of reference sequence = 99.02 ± 2.902) and variable percent identity (% identity = 52.283 ± 11.822). The second most abundant resistance gene was *AbaQ* (% identity = 72.7 ± 0.427, % length of reference sequence = 101.36 ± 0.109), a major facilitator superfamily efflux pump associated with the extrusion of quinolone-type drugs in *Acinetobacter baumannii. AbaQ* was observed across all FPES sites in 78.4% of *Pseudomonas* isolates (*Figure 6*).

**Figure 6.**
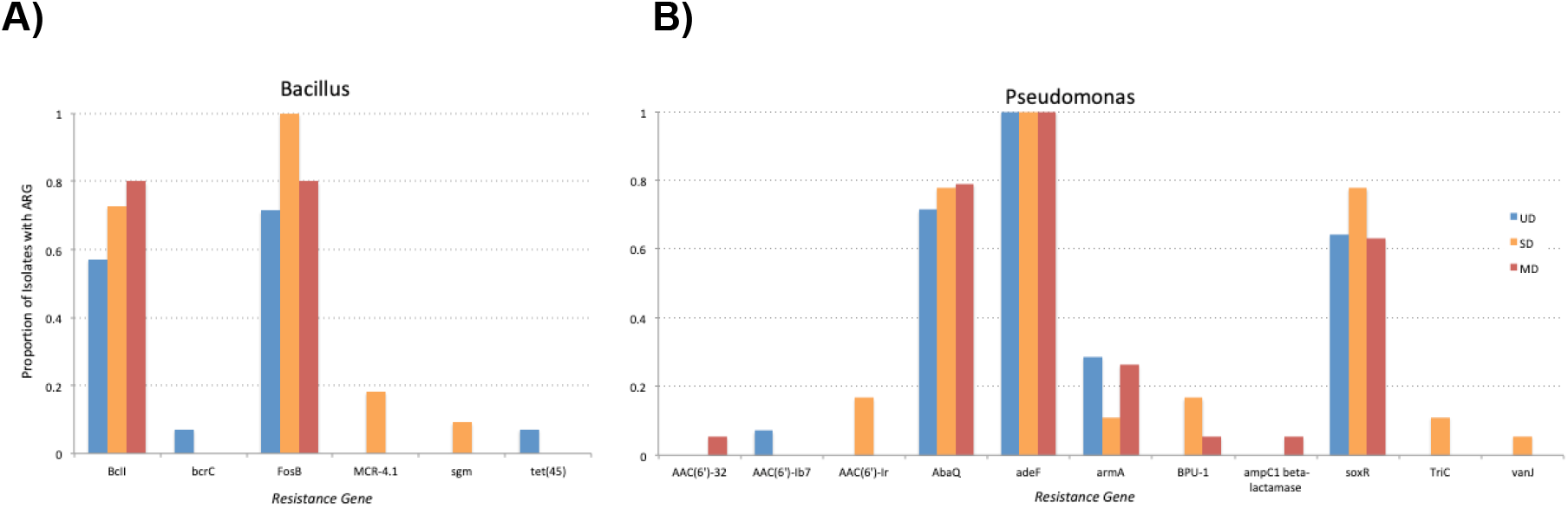
Proportion of total isolates within each FPES site carrying ARGs for A) *Bacillus* and B) *Pseudomonas* isolates.

After antibiotic efflux, genes encoding antibiotic inactivating enzymes were the most abundant found across all FPES sites and genera sampled except *Exinguobacterium*. *FosB* is a fosfomycin thiol transferase gene conferring resistance to fosfomycin, an antibiotic derived from secondary metabolites produced by soil-dwelling *Streptomyces* and *pseudomonads* that hinders cell wall synthesis by inhibiting UDP-GlcNAc enolpyruvyl transferase (Kim et al. 2012). *FosB* (% identity = *89.055 ± 2.281*, % length of reference sequence = *105.012 ± 6.992*) was the most abundant antibiotic inactivating gene with 25 chromosomally-encoded gene copies present in 83% of *Bacillus* isolates across all thaw sites. We also observed beta-lactam inactivating genes from four distinct beta-lactamase families (Table S1). Bc beta-lactamase had the highest abundance of gene copies (n=20) all encoding *BcII*, a zinc metallo-beta-lactamase that hydrolyzes a large number of penicillins and cephalosporins. *BcII* gene copies in our samples were confined to the genus *Bacillus* and found to be both highly similar (% identity = 90.755 ± 0.567) to *BcII* homologs in the CARD database and had high gene coverage (% length of reference sequence = 100.39 ± 0). Overall the high abundance of of beta-lactam resistance genes across both sites and genera directly corresponds with the high proportion of phenotypic resistance to the beta-lactam antibiotic screened, Ampicillin.

Genes encoding target alteration were the least abundant mechanism of resistance (n = 25) and included the genes armA, bcrC, MCR-4.1, Morganella morganii, gyrB, PmrF, sgm, and vanJ which are associated with resistance to aminoglycosides, peptide, glycopeptide, and fluoroquinolone antibiotics. A notable target alteration gene found was the mobilized colistin resistance (MCR) phosphoethanolamine transferase. MCR is a gene superfamily tracked by the Center for Disease Control and Prevention that confers resistance to the last resort antibiotic colistin, a critical antibiotic for treating carbapenem-resistant *Enterobacteriaceae*. We found two significant gene hits for MCR-4.1 that were fragmented on the edge of contigs and thus had low coverage (% length of reference sequence = 10.63 ± 1.174) but 100% sequence identity.

### Influence of Permafrost Thaw Distrubance

We found an increasing abundance of ARGs associated with increasing thaw distrubance (UD=101, SD=133, MD=145). We found that the mean number of CARD hits per isolate was highly significant by genus (Kruskal-Wallis p=1.9e-12; Figure 3B) and significant between the undisturbed and most-disturbed sites (Wilcoxon p = 0.045). The significant difference in ARG copies per isolate by genus suggests that the taxonomy plays a significant role in shaping abundance of resistance genes and thus with a shift in a community composition can affect the abundance and types of resistance genes. When examining the types of genes present within isolates, there is a distinct set of core ARGs found within each taxon rather than between taxa across sites such as FosB, BcII in *Bacillus* isolates and adeF, armA, and soxR in *Pseudomonas* (Figure 4). The set of intrinsic ARGs within each taxon caused groups to cluster more strongly by phylogeny rather than FPES site (Figure 4). Overall *Bacillus* isolates’ core resistance genes were comprised of primarily antibiotic inactivation genes, *Pseudomonas* was antibiotic efflux and target alteration, and *Erwinia* contained antibiotic efflux, target alteration, and inactivation. Although there were many genes found across all sites within some taxa, some genes were observed exclusively in one taxon and site such as MCR-4.1 in *Bacillus* from the semi-disturbed site. Although rare, these less abundant genes, and thus the ARGs with less of an influence on clustering, are more likely resistance determinants that were acquired either through conjugation, transformation, or transduction from members within the soil community and potentially more of an interest for assessing clinical risk.

**Figure 3.**
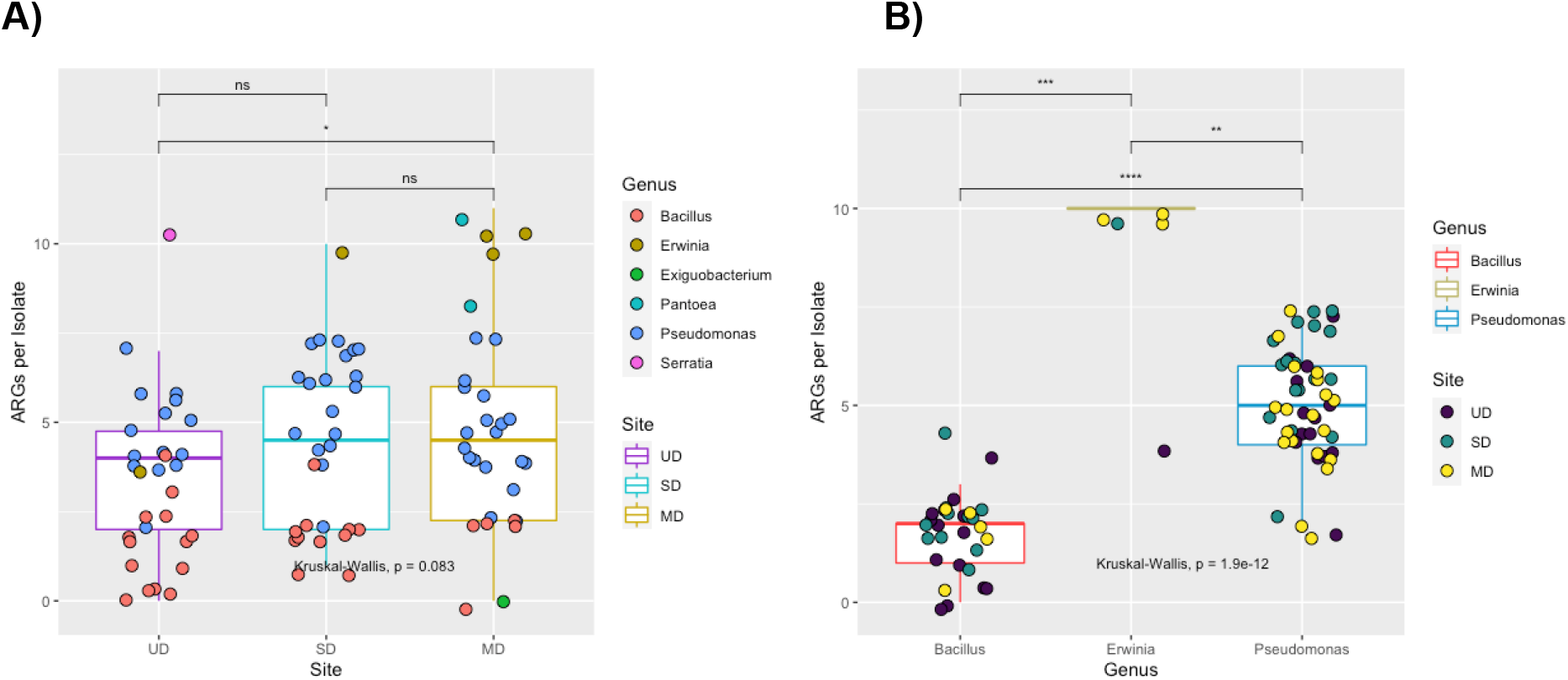
**A)** Boxplot displaying the number of antibiotic resistance genes per isolate by site. Points each represent an individual isolate color coded by genus. Wilcoxon test significance p<0.01**, <0.05*, >0.1 ns **B)** Boxplot displaying the number of antibiotic resistance genes per isolate by genus. Wilcoxon test significance p<0.01**, <0.05*, >0.1 ns Points each represent an individual isolate color coded by FPES site.

**Figure 4.**
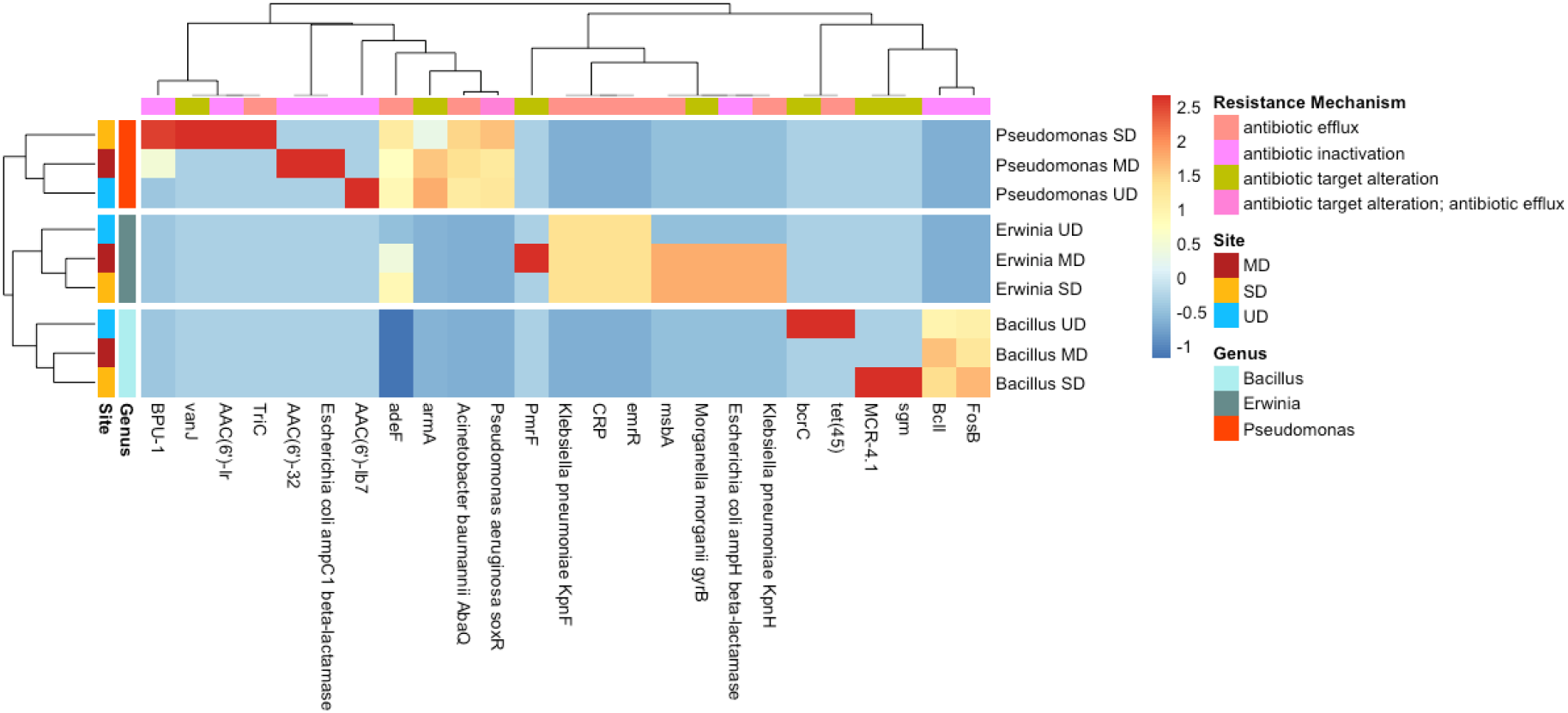
Heat map of z-score (scaled and normalized ARG count) grouped by genus and site with dendogram displaying Euclidean distance.

### RefSoil+ Comparison

Across the genera present in our FPES isolates, RefSoil+ and FPES had 15 similar ARGs, 41 unique to RefSoil+ and 12 unique to FPES (Figure 5A). The similar ARGs were primarily genes determined to be the more abundant intrinsic ARGs in FPES isolates whereas the distinct genes were often rare variants (i.e. only one copy across isolate) within each database. When comparing by genus and database we found there were significant differences in number of ARGs per isolate between *Pseudomonas* and Bacillus isolates, which was higher in FPES for *Pseudomonas* and higher in RefSoil+ for *Bacillus* (Figure 5B). Across the 90 FPES isolates, plasmids contained 4 ARO types with 6 gene copies whereas there was no significant plasmid hits across all 127 RefSoil+ isolates examined. The plasmid-borne ARGs from FPES included a BES-1 beta lactamase, two antibiotic efflux pumps (TriC and KpnF), and an undecaprenyl pyrophosphate related protein (bcrC) (Table 2).

**Figure 5.**
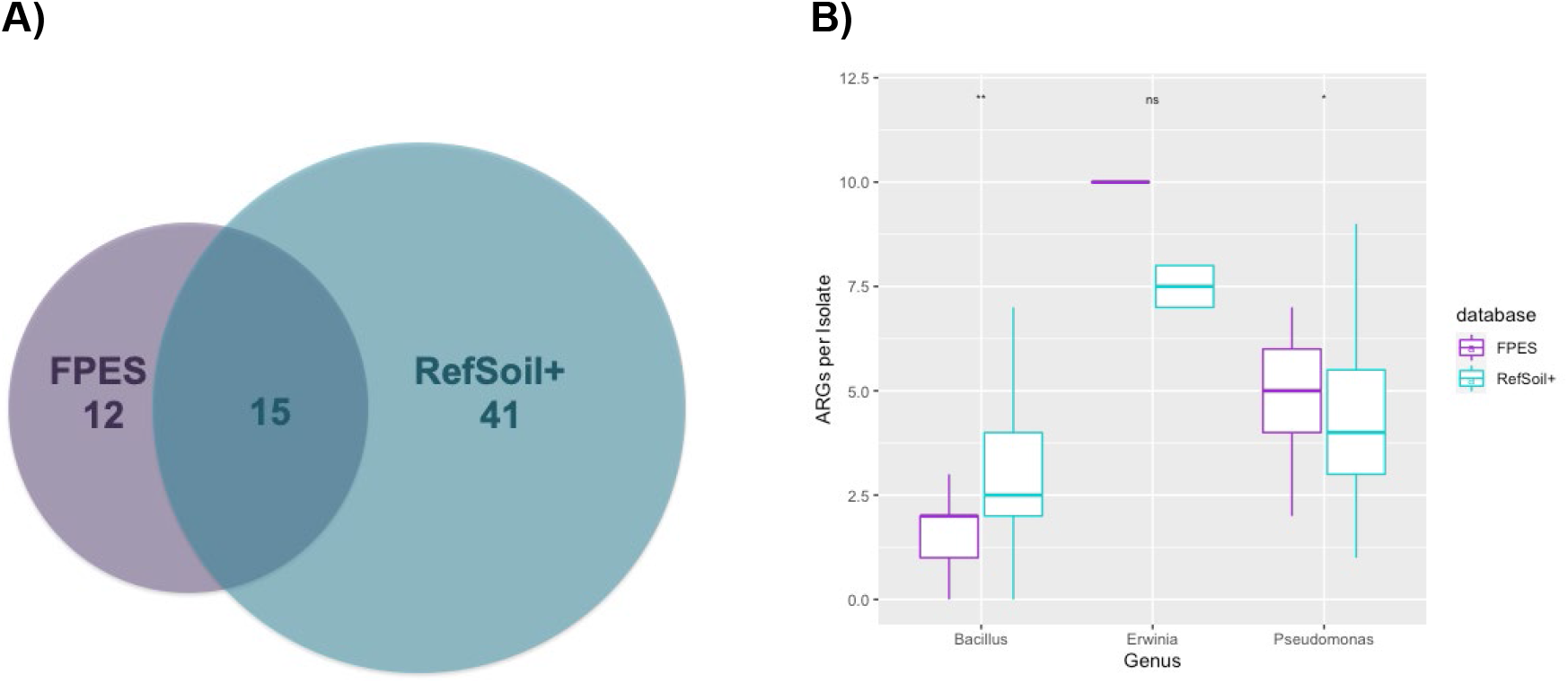
**A)** Types of ARGs by database **B)** Boxplot of the number of antibiotic resistance genes per isolate by genus and database (FPES vs RefSoil+). Wilcoxon test significance p<0.01**, <0.05*, >0.1 ns.

**Table 2.** List of ARGs and gene families found on plasmids in FPES Isolates

| Resistance Mechanism | Best_Hit_ARO | PLASMID BORNE ARGs IN FPES ISOLATES |  |  |  |
| --- | --- | --- | --- | --- | --- |
|  |  | Drug Class | AMR Gene Family | Genus origin | Count |
| antibiotic inactivation | BES-1 | carbapenem; cephalosporin; penam | SIM beta-lactamase | *Pantoea* | 1 |
| antibiotic target alteration | bcrC | peptide antibiotic | undecaprenyl pyrophosphate related proteins | *Bacillus* | 2 |
| antibiotic efflux | TriC | triclosan | RND antibiotic efflux pump | *Pseudomonas* | 2 |
|  | Klebsiella pneumoniae<br>KpnF | Broad Spectrum | MFS antibiotic efflux pump | *Erwinia* | 1 |

## DISCUSSION

The Alaskan soil bacteria in this study harbored a diverse array of resistance determinants from all major mechanisms of antibiotic resistance corroborating findings that suggest ARGs are ancient in origin and ubiquitous in soil-dwelling bacterial taxa (Wright et al. 2012, D’Costa et al. 2011). In terms of the effect of permafrost thaw, we observed a positive trend in both the proportion of resistant isolates (Figure 1) and abundance of ARGs with increasing permafrost thaw suggesting thaw plays a role in shaping the resistome. This is further supported by our findings that showed a significant difference in the mean number of ARGs per genome between the undisturbed and most-disurbed sites (Figure 3A). However, the mean ARGs per genome by genus (p= 1.9e-12) was much more highly significant than by FPES site (p= 0.045) suggesting the positive trend in abundance of ARGs across FPES sites is a result of the increasing number of randomly sampled Proteobacteria (*Erwinia, Pseudomonas, Pantoea*, and *Serratia*) with thaw (n= 24 MD; 19 SD;15 UD). Isolates from *Proteobacterial* phyla had a significantly higher number of ARGs per genome compared to the Firmicutes (*Bacillus* and *Exiguobacterium*) (Figure 3B). The link between host taxa and ARG abundance within this cultivatable subset of the active layer bacterial community suggests an increasing abundance of Proteobacteria with thaw can affect the abundance of resistance genes comprising the resistome.

Along with the observed association of host taxa and ARG abundance, we found that the ARGs clustered more strongly by bacterial genera than thaw site (Figure 4) and that the predominant mechanism of resistance (e.g. efflux, inactivation, target protection, and target alteration) was dependent on host taxa. This connection between host taxa and resistance determinants means that as microbial community composition shifts in response to permafrost thaw (Schurr et al. 2007), so will the predominant taxa shaping the types, abundance, and mobility of ARGs within the resistome. Proteobacterial genera predominately harbored ARGs encoding efflux pumps (mean = 4.93 per isolate) and very few encoding antibiotic inactivating enzymes (mean = 0.28 per isolate) whereas the most abundant resistance mechanism in *Bacillus* spp. was antibiotic inactivation (mean = 1.5 per isolate) and the one of the lowest abundance mechanisms was ARGs encoding efflux pumps (mean = 0.03 per isolate). Within each bacterial genus there was a distinct set of core ARGs that were chromosomally encoded and ubiquitous across thaw levels within a genus, such as *adeF* in *Pseudomonas* and *BcII* in *Bacillus*. These core genes are usually less prone to horizontal gene transfer compared to accessory determinants associated with genomic hotspots and mobile genetic elements such as integrons, plasmids, transposons (Oliveira et al. 2017; Harrison & Brockhurst 2012). Yet core resistance genes in soil bacteria still pose a risk because they are widespread within taxa as an intrinsic part of the genome that likely plays a role in both the colonization of the rhizosphere, which contains toxic compounds from plants and associated microbiota, and a high-level antibiotic resistance found with many environmental borne opportunistic pathogens such as *Pseudomonas aeruginosa* (Martinez 2009, Gallanger et al. 2017).

Some of the isolates in this study do in fact belong to taxa of known opportunistic human pathogens, such as *Pantonea agglomerans, Bacillus cereus*, and several *Pseudomonas spp*. and were found to carry both chromosomally encoded intrinsic resistance determinants and plasmid-borne ARGs (Kotiranta et al. 2000, Cruz et al. 2007). However, even non-pathogenic soil bacteria regularly interact with waterways, air, and built habitats, such as hospital surfaces, and can carry resistance determinants within the genome that can be disseminated to bacteria of clinical importance via horizontal gene transfer. A study by Hu et al. 2016 analyzed the mobilome of 23,425 bacterial genomes and found that mobile ARGs are mainly present in four bacterial phyla, the top two of which were *Proteobacteria* (399 mobile ARGs) and *Firmicutes* (86 mobile ARGs). All of the FPES isolates belong to these two phyla and twelve isolates were shown to carry plasmid-borne ARGs. Although only 1.6% of the ARGs copies from FPES isolates were located on plasmids, presence on these MGEs is telling of the low, but real, potential for ARGs from these soils to be disseminated. Moreover the higher proportion of plasmid-borne ARGs in FPES isolates compared to the RefSoil+ genomes of bacteria from the same genera, suggests the role local soil attributes have in selecting for plasmid-carriage further highlighting the clinical significance of ARGs harbored in Alaskan soils.

In our soil isolates from across FPES sites, resistance-nodulation-cell division (RND) efflux pumps were the most abundant antibiotic resistance family found in all *Proteobacteria*. RND efflux pumps have been described as a major tolerance mechanism allowing the effective extrusion of organic solvent from the interior of the cell to the exterior environment, this mechanism is found to be especially prevalent in *Pseudomonas* (Ramos et al. 2001). As environmental change increasingly affects the arctic in the form of higher annual ambient air temperature and anthropogenic disturbance of soils, microbial life within the active layer have to cope with the release of biogenic volatile organic compounds (BVOC) amplified by thawing permafrost (Kramshøj et al. 2018). Efflux pumps therefore provide an effective mechanism for coping with toxic substances, such as antibiotics, and changes of BVOC concentration within active layer soils associated with climate change.

Although this study does not provide direct evidence of a horizontal gene transfer event from soil bacteria to pathogenic bacteria, it identifies the remarkable abundance and diversity of antibiotic resistance in Alaskan soils. The resistance genes identified here are a restricted representation of the Alaskan soil community because this study is limited in geography, to the CARD ARG database, and to the culturable non-fastidious aerobic to facultative anaerobic bacteria of the phyla *Proteobacteria* and *Firmicutes*. Despite this restricted scope, the remarkable abundance, diversity, and presence on plasmids is suggestive of the extent these soils have as a reservoir for antibiotic resistance and for the potential to compromise health.

## CONCLUSION

As antibiotic resistance continues to emerge and rapidly spread in clinical settings, it is imperative to generate studies that build insight into the ecology of environmental resistance genes that pose a threat to human health. This study provides insight into the occurrence of diverse ARGs found in Alaskan soil bacteria which is suggestive of the potential to compromise health. The observed differences in ARG abundance with increasing permafrost thaw suggest the role of soil disturbance in driving the distribution of resistant determinants and the predominant phyla that shape the resistome, such as adeF efflux pump genes in *Proteobacteria* which can provide effective tolerance mechanism for dealing with change in thawing Alaskan permafrost. Moreover, the high-quality whole genome assemblies generated in this study are an extensive resource for microbial researchers interested in permafrost thaw and will provide a steppingstone for future research into ARG mobility and transmission risks.

## DATA AVAILABILITY

This genome project is indexed at GenBank under BioProject accession number XXXX. These whole genomes have been deposited in GenBank under the accession nos. XXXX-XXX. Raw sequencing data for this project can be found in the GenBank SRA under XXXX.

## ACKNOWLEDMENTS

Many thanks go to Ursel Schütte, Taylor Seitz, Scout McDougal, Anne-Lise Ducluzeau, Jennie Humphrey and Sara Kline for laboratory help. Tom Douglas from the Cold Regions Research and Engineering Laboratory (CRREL) Alaska who provided field site access.

We acknowledge generous support from the Institute of Arctic Biology, Alaska INBRE, and the BLaST program. Research reported here was supported by BLaST through the National Institute of General Medical Sciences of the National Institutes of Health under awards UL1GM118991, TL4GM118992, and RL5GM118990. Research reported in this publication was supported by an Institutional Development Award (IDeA) from the National Institute of General Medical Sciences of the National Institutes of Health under grant P20GM103395.

